# The freeze-dried extracts of *Salvia coccinea* Juss. Ex Murray attenuate myocardial ischemia reperfusion injury in a global ischemia Rat model

**DOI:** 10.1101/396119

**Authors:** Nelly Murugi Nyaga, Peter Waweru Mwangi, Frederick Bukachi

## Abstract

**Background:** Ischemia reperfusion injury is the leading cause of myocardial cell death in Ischemic Heart Disease. Thus intensive research efforts are geared at discovering pharmacological approaches that prevent it. Over twenty species from the genus *Salvia* are widely applied in traditional Chinese medicine in the management of heart diseases with *Salvia miltiorrhiza* (Danshen) being a canonical example. Our study aimed to investigate the cardio-protective effects of the freeze-dried extracts of *salvia coccinea* against ischemia reperfusion injury in a rodent *in-vitro* model of global ischemia.

**Methods:** Forty two (42) Sprague Dawley rats were randomly assigned into five groups: positive control (Glucosamine 1000mg/kg), negative control group (Krebs Henseleit buffer), low dose test (50 mg/100ml), medium dose test (100 mg/100ml), and high dose test (200 mg/100ml).

The cardio-protective effects of the different treatments were evaluated in a global ischemia model using isolated rat hearts mounted on a Langendorff system.

Naloxone 2.2 μmol/L (μ opioid receptor blocker), and theophylline 1000 μmol/L (non-specific adenosine receptor blocker) were co-administered with 50 mg of *S.coccinea* in the mechanism of action experiments.

The following indices of cardiac function were recorded pre- and post-ischemia: left ventricular developed pressure (LVDP), heart rate, and maximum rate of contraction and relaxation. All data were expressed as Mean ± Standard Error of Mean and analyzed using one-way ANOVA and Tukey post-hoc tests. Significance was set at p < 0.05.

**Results:** The freeze-dried extracts of *S. coccinea* had significant effects on post-ischemic contractile function recovery in the early [51.4 ± 9.7% (low dose test) vs. 14.9 ± 3.3% (medium dose test) vs. 12.7 ± 2.6% (high dose test) vs. 13.7 ± 5.7% (negative control): p<0.05] and late [38.6 ± 8.9% (low dose test) vs. 22.0± 7.1% (medium dose test) vs. 14.6 ± 5.8 (high dose test) vs. 12.5 ± 4.2% (negative control): p< 0.05]. Reperfusion phases with the highest LVDP recovery were observed at the 50 mg dosage level.

The freeze-dried extracts of *S. coccinea* had significant negative chronotropic effects on heart rate [234.0 ± 2.4 beats/min to 90.0 ± 7.0 beats/min, 50 mg vs. 102.0 ± 13.9 beats/min to 135.0 ± 25.9 beats/min, control P<0.05].

The cardioprotective effects of *S. coccinea* displayed an inverted U-shaped dose-response curve with low dose stimulation and high dose inhibition.

Naloxone completely abolished the LVDP recovery afforded by the freeze-dried extracts of *S. coccinea* at the 50 mg dosage level while adenosine only partly abolished the LVDP recovery (9.5 ± 3.2% (naloxone) *vs.* 15.5 ± 5.8% (adenosine): P>0.05).

**Conclusion:** The freeze-dried extracts of *S. coccinea* possessed significant cardioprotective effects which appear to be mediated by activation of the opioidergic pathway in the heart.

## 1.0 Introduction

Cardiovascular diseases are the leading cause of death globally with a 31% annual death rate (1). Ischemic heart disease (IHD) is responsible for more than 40% of these cardiovascular deaths and thus poses a major global health problem(2).

The clinical manifestations of IHD result from subjection of the heart to ischemia-reperfusion injury which leads to myocardial injury, cardiac dysfunction, arrhythmias, heart failure and ultimately death (3).

Cardioprotection aims to prevent the death of cells during acute injury and may be achieved through myocardial or pharmacological conditioning carried out pre- or post-ischemia (4).

Salvia species are used widely in Traditional Chinese Medicine (TCM) as Danshen to manage cardiac conditions. *Salvia coccinea* has been used since the 17^th^ century by the Chinese who refer to it as “cure all” or “sage the savior” for the management of common maladies (5). In South America a boiled mixture is used to bathe varicosities and treat blood clots (6). The aqueous and alcohol extracts of *Salvia coccinea* contain potent phenolic antioxidants (7). The present study aimed to investigate the cardioprotective effects of the freeze-dried extract of *S. coccinea* against ischemia reperfusion injury and to determine its mode of action.

## 2.0 Materials and methods

### 2.1 Plant material and freeze dried extract preparation

The specimen of *Salvia coccinea* was harvested from Naivasha County, Kenya, and its identity confirmed by resident taxonomists of the herbarium situated at the Department of Botany, School of Biological sciences, University of Nairobi.

The whole plant was used in the study. The plant parts were cut into small pieces and air dried for one week. The dried plant was milled using a standard mill and the resulting powder macerated in cold distilled water in a ratio of 1:10 (weight: volume) with vigorous shaking for an hour. The suspension obtained was filtered consecutively using cotton wool and filter paper (Whatman’s^TM^). The filtrates were frozen and lyophilized and the resultant freeze-dried extracts weighed and placed in sample containers. These were stored in a standard laboratory freezer (Hotpoint™). During each procedure, the freeze-dried extracts were reconstituted to form the required concentrations.

### 2.2 Experimental Animals

Forty-two (42) six month old adult male Sprague Dawley rats, weighing 300-400 g were used. The animals were housed in standard cages containing five rats in each. The experiments were performed in the animal house at the Department of Medical Physiology, University of Nairobi. The following conditions were maintained in the animal house: constant room temperature (23 +/− 2°C) and 30-70% relative humidity with a 12-hr. light/day cycle. The animals were provided with standard rat chow pellets and tap water ad libitum. The protocol was approved by the Biosafety, Animal Use and Ethics Committee, Faculty of Veterinary Medicine, University of Nairobi. Ref. Number: (FVM BAUEC/2016/113). All experiments were based on internationally accepted directions for the use and care of laboratory animals as stipulated by the FELASA guidelines (8).

### 2.3 Drugs

Naloxone HCL (Sanofi Ltd.), Theophylline (GlaxoSmithKline Ltd.), Glucosamine (Power Health Ltd.), and Heparin (Rotex Medica Ltd.) were obtained from Dischem Pharmacy, Nairobi. Sodium pentobarbital (Bayer Ltd.) was obtained from Lesukut Ltd. NaCI, KCl, KH_2_PO4, MgSO_4_, NaHCO_3_, CaCl_2_, and glucose (LOBA Ltd.) were obtained from Chem-labs, Nairobi. All drugs were freshly prepared in normal saline on the day of the experiment.

### 2.4 Isolated heart preparation

Each rat was anaesthetized with sodium pentobarbital 6% (40 mg/kg) intraperitoneally. There after heparin (500 u/kg) was administered intravenously immediately the rat lost its righting reflex. The hearts were excised through a trans-abdominal incision and immediately immersed in ice-cold Krebs’ Henseleit (KH) buffer solution. A plastic cannula was inserted through the aorta, clamped with a blunt artery clip and retrograde perfusion with Krebs Henseleit buffer (10 ml/min) initiated.

### 2.5 Measurement of isovolumic cardiac performance

A latex balloon connected to a pressure transducer was inserted into the left ventricle via the mitral valve and inflated to increase the diastolic pressure by regulating the volume of saline in it. The pressure transducer was connected to a power lab (AD Instruments, Colorado Springs, CO, USA) recording system. The following cardiac indices were then measured and averaged from twelve (12) beats: Left ventricular developed pressure (LVDP), maximum rate of contraction (+dp/dt_max_), maximum rate of relaxation (−dp/dt_min_), and the heart rate.

### 2.6 Cardioprotection experiment

The hearts were equilibrated for twenty (20) minutes and subjected to thirty (30) minutes global ischemia by halting perfusion and immersing the hearts in a KH buffer filled organ bath. The hearts were divided into five (5) groups and on reperfusion treated (Table 1).

**Table 1:** Heart treatment during reperfusion

| **TREATMENT**<br>**STEPS**<br>**Time**<br>**(minutes)** | **EXPERIMENTAL GROUPS** |  |  |  |  |
| --- | --- | --- | --- | --- | --- |
|  | **POSITIVE**<br>**CONTROL** | **CONTROL** | **50 mg** | **100 mg** | **200 mg** |
| 15 | Equilibration | Equilibration | Equilibration | Equilibration | Equilibration |
| 30 | Global<br>Ischemia | Global<br>Ischemia | Global<br>Ischemia | Global<br>Ischemia | Global<br>Ischemia |
| 90 | Reperfusion<br>with KH<br>buffer +<br>Glucosamine | Reperfusion<br>with KH<br>buffer<br>containing<br>*Salvia*<br>*coccinea*<br>extract (50<br>mg per liter) | Reperfusion<br>with KH<br>buffer<br>containing<br>*Salvia*<br>*coccinea*<br>extract (100<br>mg per liter) | Reperfusion<br>with KH<br>buffer<br>containing<br>*Salvia*<br>*coccinea*<br>extract (200<br>mg per liter) | Reperfusion<br>with<br>plain KH<br>buffer |

*2.7 Mechanism of action experiments*

### 2.7 Mechanism of action experiments

Theophylline and Naloxone, which are non-selective adenosine and opioid receptor blockers respectively, were used to determine the signaling pathway of cardioprotection by *Salvia coccinea*. These blockers were administered for 20 minutes at the onset of reperfusion before treatment with *Salvia coccinea* extracts (Table 2).

**Table 2:** Heart treatment during reperfusion

| TREATMENT<br>STEPS<br>Time<br>(minutes) | EXPERIMENTAL GROUPS |  |  |  |  |
| --- | --- | --- | --- | --- | --- |
|  | POSITIVE<br>CONTROL | CONTROL | 50 mg | 100 mg | 200 mg |
| 15 | Equilibration | Equilibration | Equilibration | Equilibration | Equilibration |
| 30 | Global<br>Ischemia | Global<br>Ischemia | Global<br>Ischemia | Global<br>Ischemia | Global<br>Ischemia |
| 90 | Reperfusion<br>with KH<br>buffer +<br>Glucosamine | Reperfusion<br>with KH<br>buffer<br>containing<br>*Salvia*<br>*coccinea*<br>extract (50<br>mg per liter) | Reperfusion<br>with KH<br>buffer<br>containing<br>*Salvia*<br>*coccinea*<br>extract (100<br>mg per liter) | Reperfusion<br>with KH<br>buffer<br>containing<br>*Salvia*<br>*coccinea*<br>extract (200<br>mg per liter) | Reperfusion<br>with<br>plain KH<br>buffer |

### 2.9 Data and statistical analyses

The experimental data were expressed as the mean +/−SEM. The data were analyzed using One-way ANOVA with post-hoc statistical analysis being performed using Tukey HSD in cases of significance which was set at p<0.05. The Kruskall Wallis non-parametric with Dunns’ post-hoc statistical test in cases of significance (p<0.05) was used to analyze the extent of the histological damage. GraphPad Prism 6 suite of statistical software was used for carrying out the statistical tests.

## 3.0 Results

### 3.1 Effects on cardiac function

#### 3.1.1 Baseline cardiac function

There were no significant differences in LVDP between the groups during equilibration [43.8 ± 2.5 mmHg (positive control) vs. 40.9 ± 7.5 mmHg (negative control) vs. 51.1 ± 1.4 mmHg (low dose test) vs. 54.8 ± 1.3 mmHg (medium dose test) vs. 40.6 ± 2.6 mmHg (high dose test); P=0.05]. The graphical representation of the equilibration LVDP values is shown in Figure 1.

**Figure 1.**
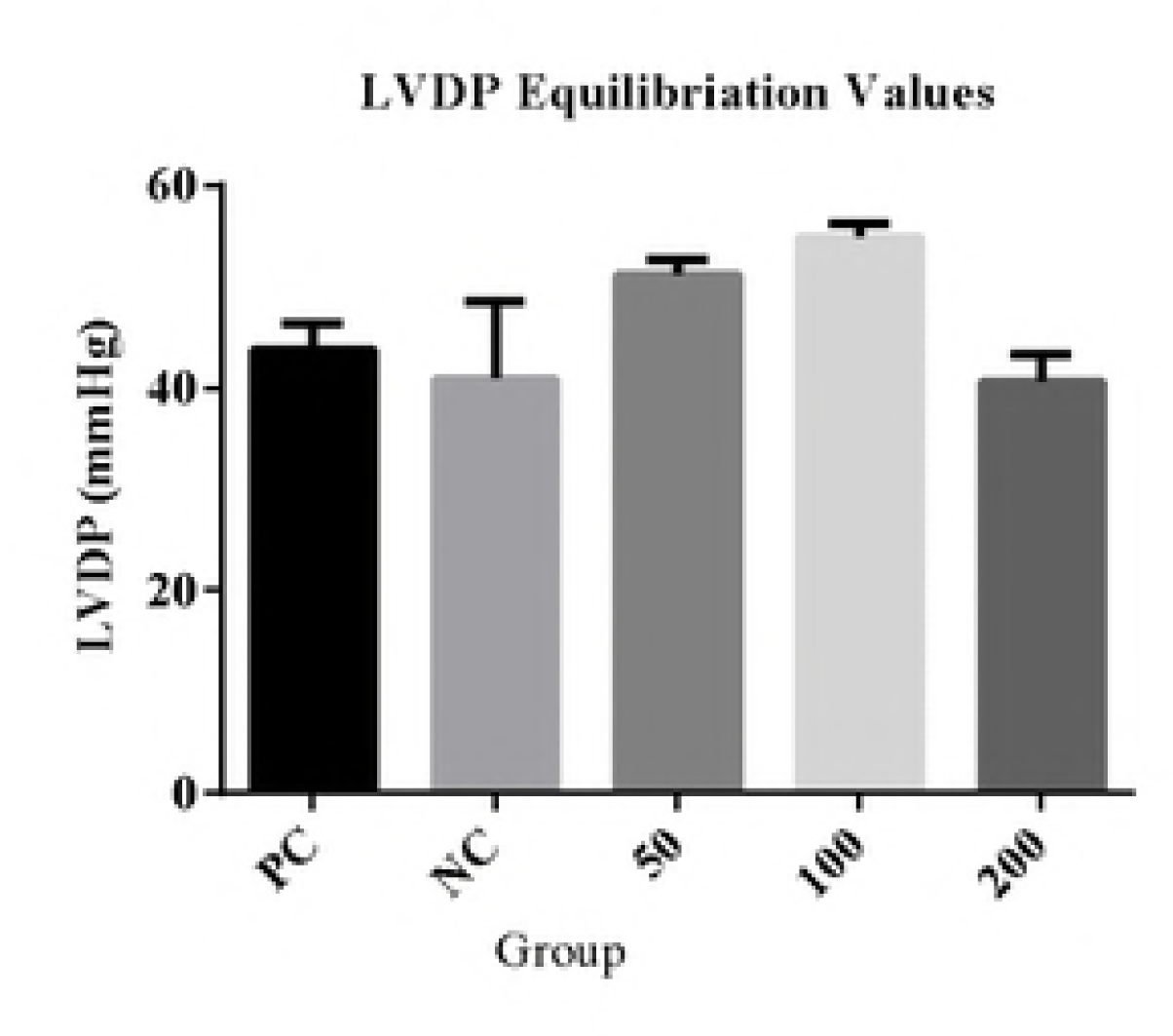
Equilibration Left Ventricular Developed Pressure (LVDP) for each group. PC (positive control), NC (Negative control).

#### 3.1.2 Effects of S. coccinea extracts on the left ventricular developed pressure (LVDP)

The freeze-dried extracts of *Salvia coccinea* had significant effects on the LVDP recovery both in the early [51.4 ± 9.7% (low dose test) vs. 14.9 ± 3.3% (medium dose test) vs. 12.7 ± 2.6% (high dose test) vs. 13.7 ± 5.7% (negative control): p<0.05] and late [38.6 ± 8.9% (low dose test) vs. 22.0± 7.1% (medium dose test) vs. 14.6 ± 5.8 (high dose test) vs. 12.5 ± 4.2% (negative control): p< 0.05] phases of reperfusion. Post-hoc statistical analysis using Tukey’s multiple comparison test showed that the low dose (50 mg) had a significantly higher cardioprotective activity in the early phase compared to the medium dose (100 mg), high dose (200 mg) and negative control (KH) groups (p<0.05). The 50 mg dose also had significant cardioprotective activity in the late phase compared to the KH and 200 mg group (p<0.01). A graphical representation of the LVDP percentage recovery is shown in Figure 2.

**Figure 2.**
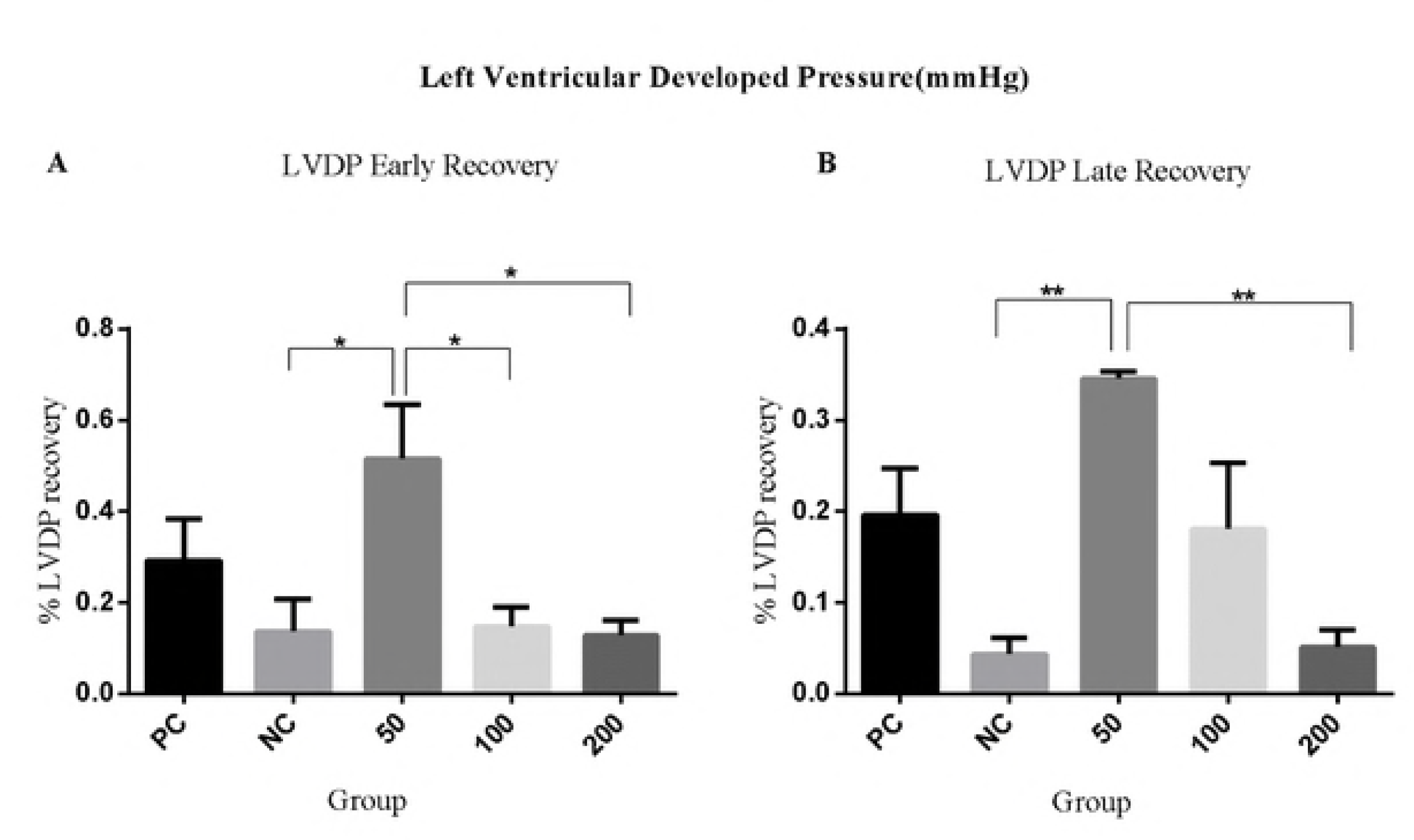
Post ischemic Left Ventricular Developed Pressure. A) Percentage early recovery after post ischemic post conditioning with *S. coccinea*. B) Percentage late recovery after post ischemic post conditioning with *S. coccinea*. LVDP (Left Ventricular Developed Pressure), PC (positive control), NC (Negative control). KEY: *= P ≤ 0.05; **P ≤ 0.01

#### 3.1.3 Effects of the S. coccinea extracts on the maximum rate of contraction (+dp/dt max)

The freeze-dried extracts of *S.coccinea* had a significant effect on the recovery of the +dp/dt _max_ in the early [78.2 ± 3.8% (low dose test) vs. 18.7 ± 3.9% (medium dose test) vs. 37.9 ± 8.1% (negative control) vs. 41.4 ± 8.9% (high dose test): P<0.05] and late phases [(69.8 ± 6.9% (low dose test) vs. 43.5 ± 6.6% (medium dose test) vs. 40.2 ± 14.6% (high dose test) vs. 44.8 ± 2.5% (negative control): p<0.05] of reperfusion. Post-hoc statistical analysis using Tukey’s multiple comparison test showed that the extract at 50 mg dose had significant cardioprotective activity in the early phase compared to the glucosamine group (P < 0.05) and 100 mg group (P<0.01). A graphical representation of the +dp/dt _max_ percentage recovery is shown in Figure 3.

**Figure 3.**
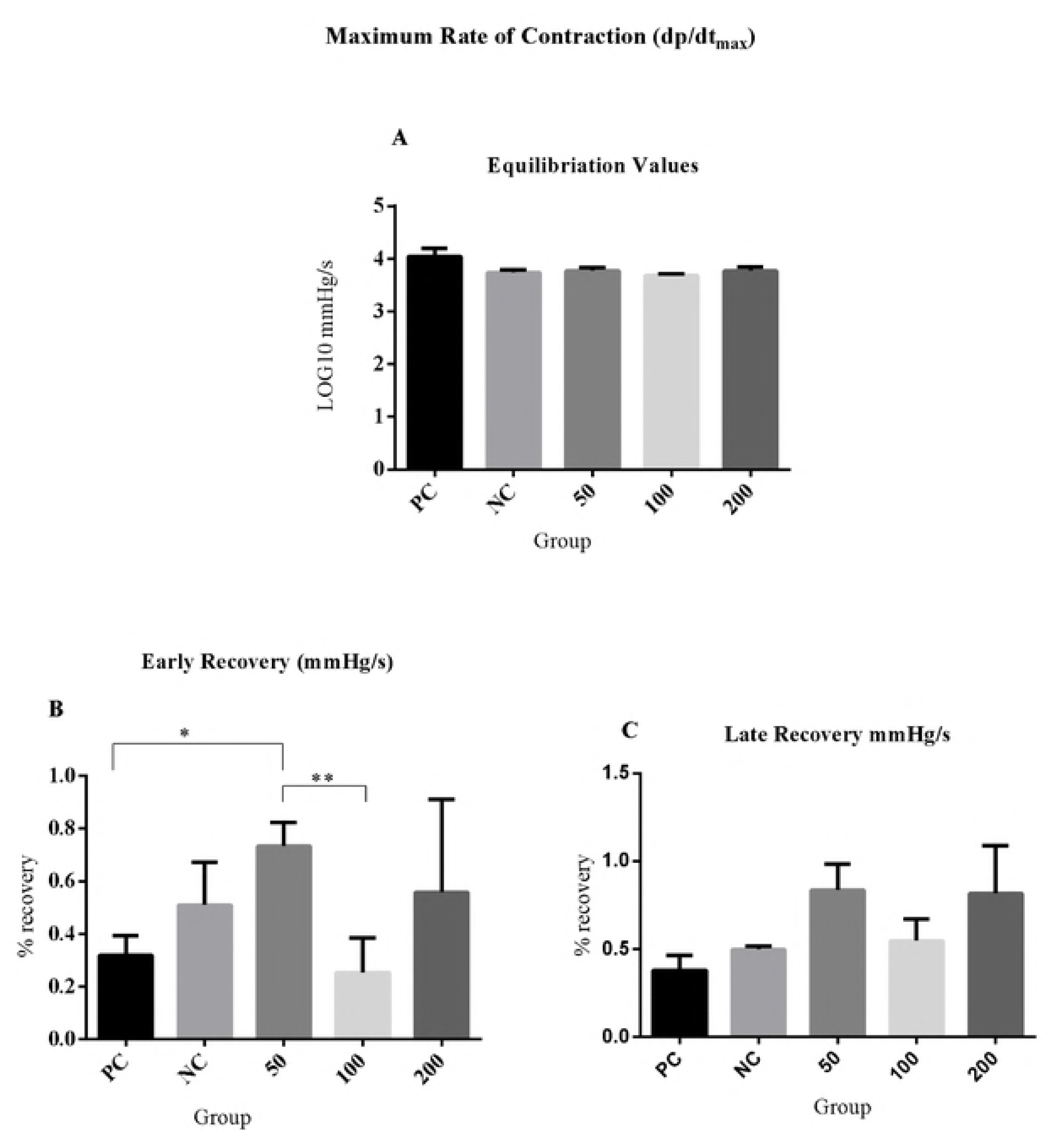
Post ischemic calculated maximum rate of contraction. A) Equilibration values B) Percentage early recovery after treatment with *s. coccinea* C) Late recovery after treatment with *s. coccinea.* LVDP (Left Ventricular Developed Pressure), PC (positive control), NC (Negative control).KEY: *=P ≤ 0.05; **=P ≤ 0.01.

#### 3.1.4 Effects of the S. coccinea extracts on the maximum rate of relaxation (−dp/dtmin)

The freeze-dried extracts of *Salvia coccinea* caused a significant −dp/dt _min_ recovery in the early [131.6 ± 8.3 (low dose test) vs. 45.6 ± 5.3% (medium dose test) vs. 55.2 ± 13.1% (high dose test) vs. 83.8 ± 11.8% (negative control): P<0.05] and late phases [149.0 ± 23.7% (low dose test) vs. 97.4 ± 10.8% (medium dose test) vs. 78.0 ± 31.0% (high dose test) vs. 113.6 ± 4.6% (negative control): P<0.05] of reperfusion. Post-hoc statistical analysis using Tukey’s multiple comparison test showed that the low dose test (50 mg) possessed significant cardioprotective activity in the early phase compared to the negative control (KH) group (P <0.01), the medium dose test (100 mg) group (P<0.001), the high dose test (200 mg) group (p<0.05) and the test positive control (glucosamine) group (P<0.01). The low dose also had significant cardioprotective activity in the late phase compared to the negative control (P <0.05), the high dose test (p<0.01), the medium dose test (P<0.05) and the positive control (P < 0.01) A graphical representation of the −dp/dt _min_ percentage recovery is shown in Figure 4.

**Figure 4.**
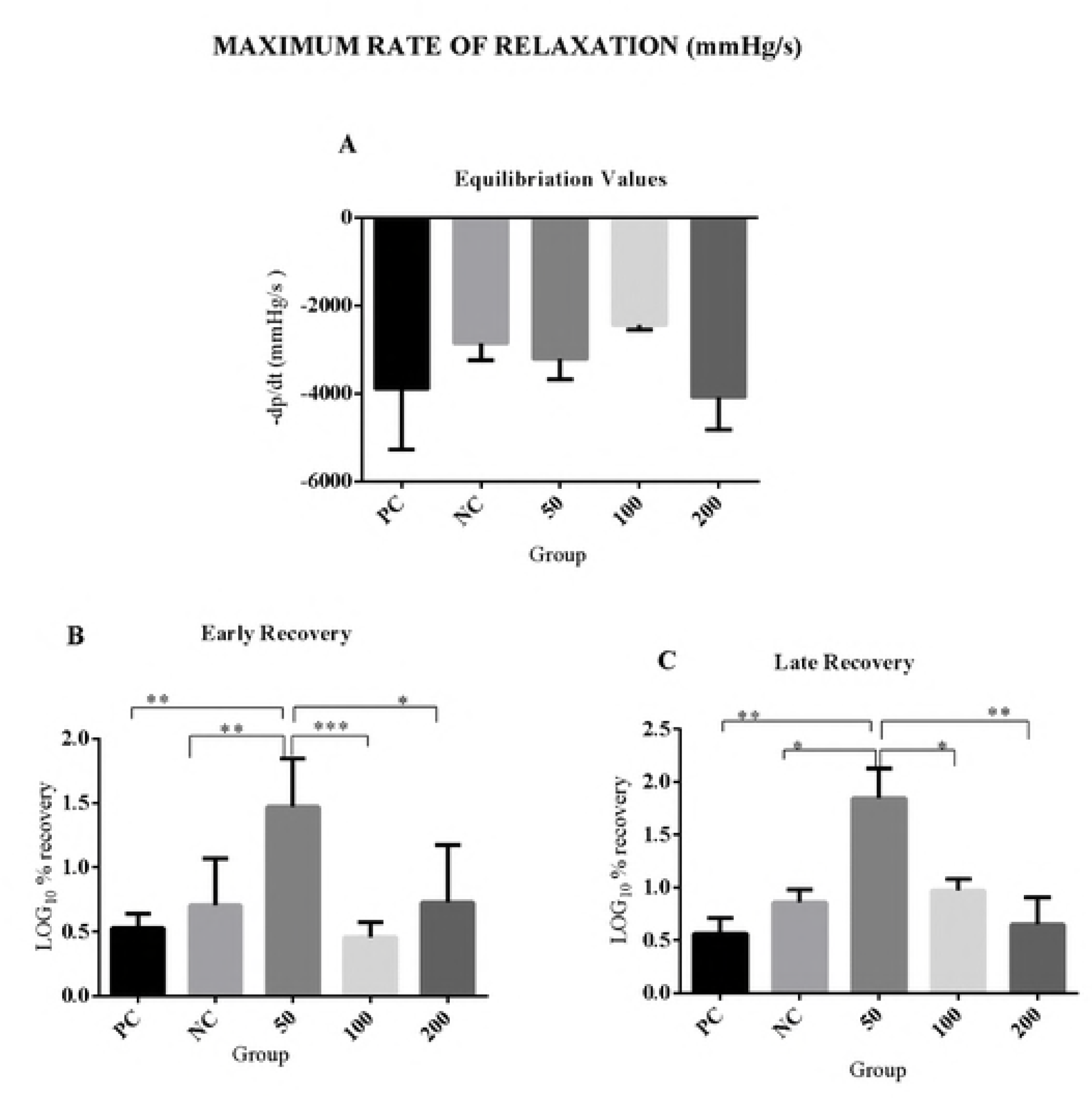
Post ischemic calculated maximum rate of relaxation. A) Equilibration values B) Percentage early recovery after treatment with *S. coccinea* C) percentage late recovery after treatment with *S. coccinea*. LVDP (Left Ventricular Developed Pressure), PC (positive control), NC (Negative control).KEY: *=P≤0.05; **=P≤0.01; ***=P≤ 0.001.

#### 3.1.5 Trends of recovery in the early and late phases of reperfusion

The trend of recovery was obtained by plotting the ratios of the percentage recovery in both phases for each of the cardiac indices to determine the effectiveness of *Salvia coccinea* in early and late reperfusion. The low dose test group had the highest recovery rate in both phases. The graphical representation of these trends is shown in figure 5, 6 and 7.

**Figure 5.**
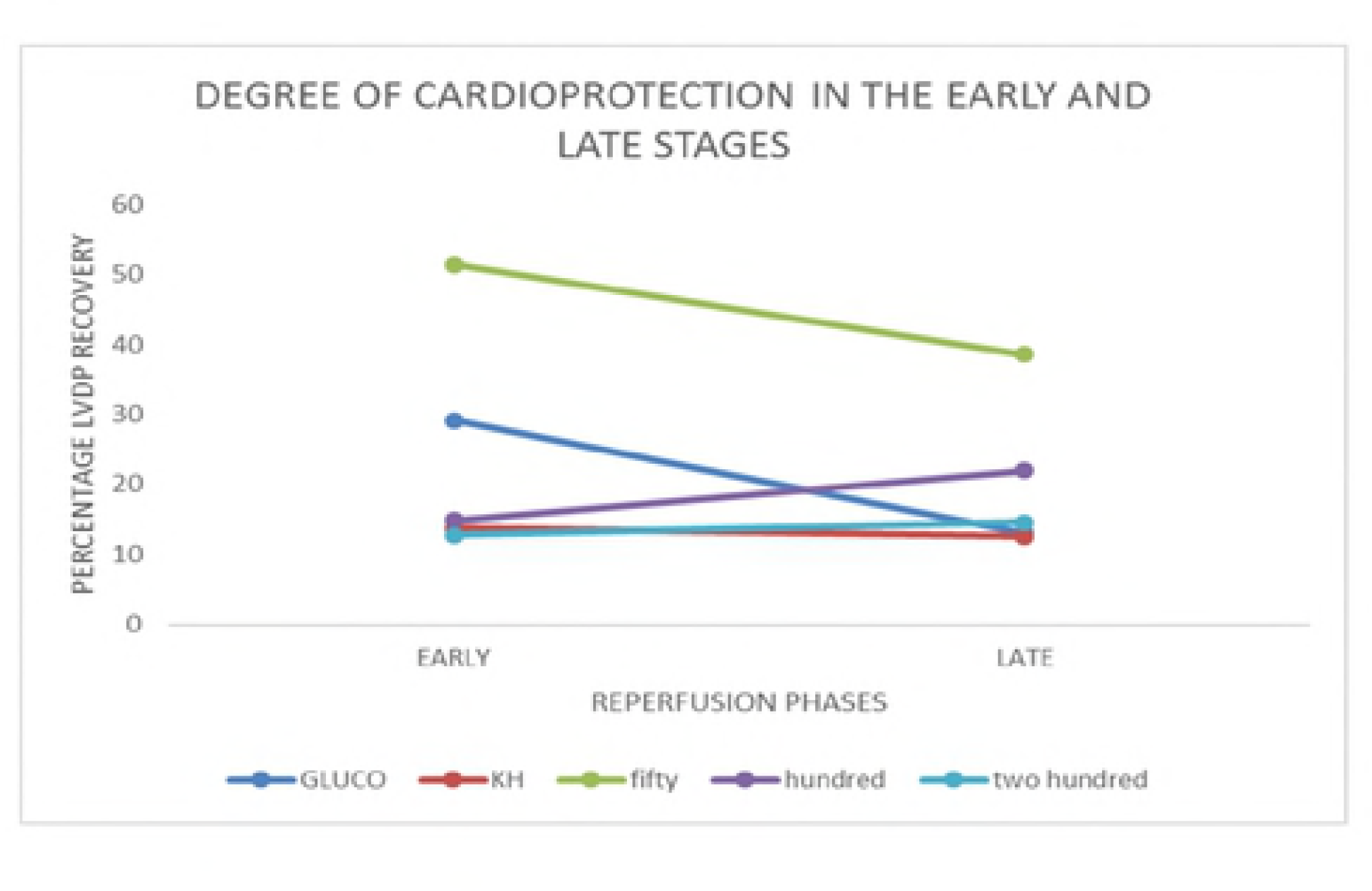
Trends of LVDP recovery in the early and late reperfusion phases.

**Figure 6.**
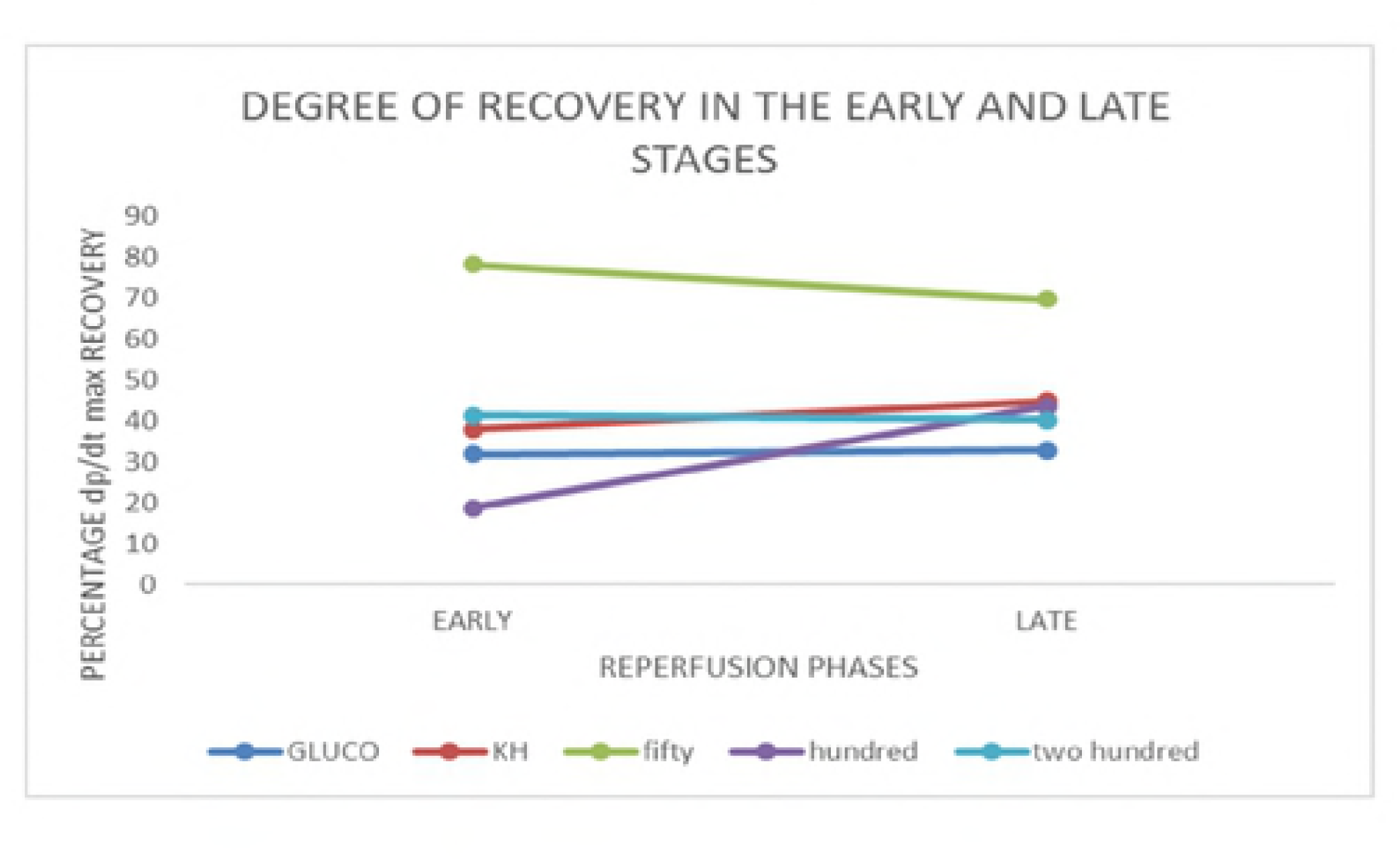
Percentage +dp/dt _max_ recovery in the early and late reperfusion phases.

**Figure 7.**
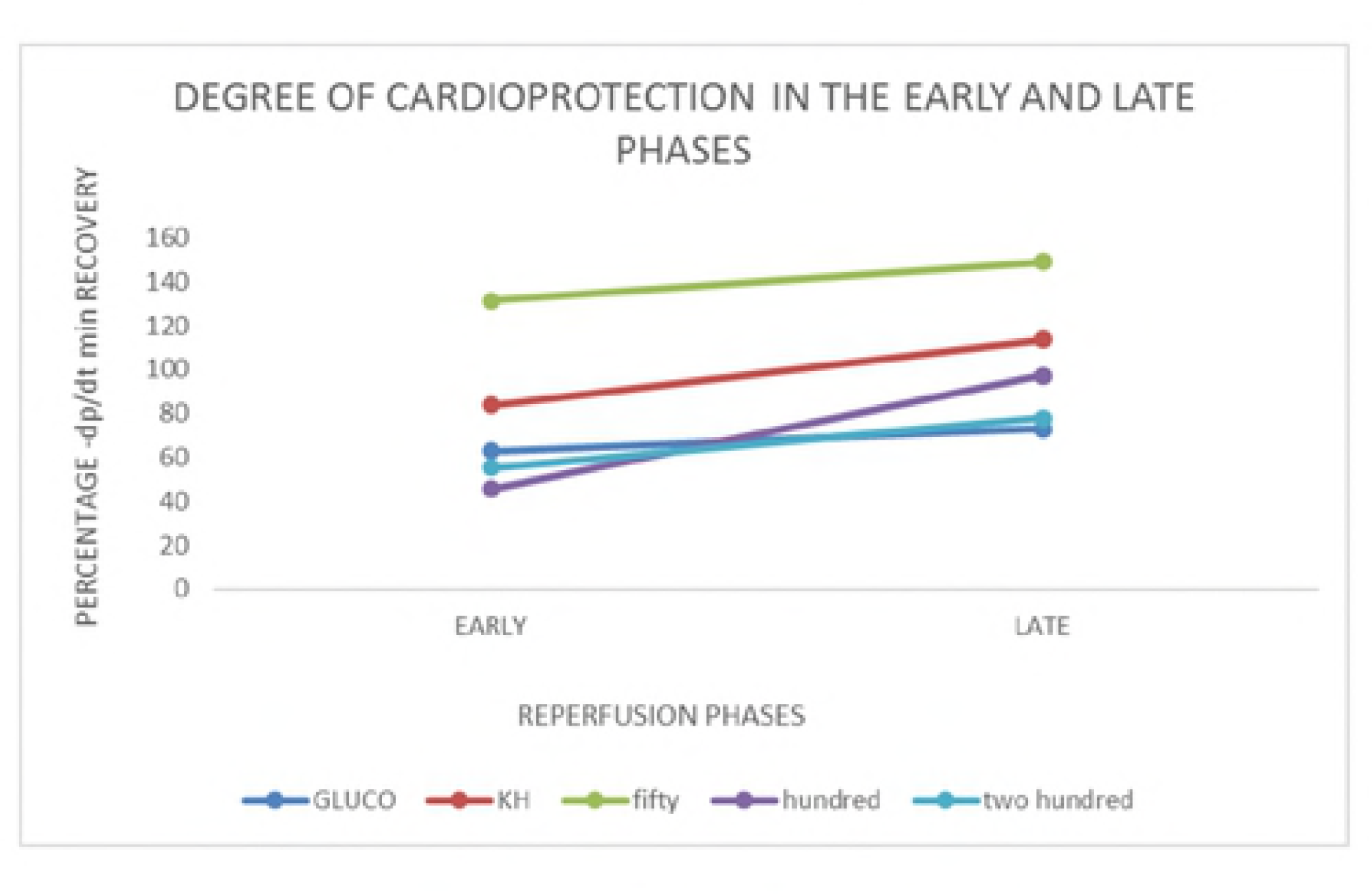
Trends of −dp/dt _min_ recovery in the early and late reperfusion phases.

#### 3.1.6 Effects of the freeze-dried extracts of S. coccinea on heart rate

The freeze-dried extracts of *S.coccinea* caused a significant decrease in heart rate [234.0 ± 2.4 beats/min (low dose test) vs. 258.0 ± 11.5 beats/min (medium dose test) vs. 174.0 ± 32.6 beats/min (high dose test)] in the early [90.0 ± 7.0 beats/min (low dose test) vs. 98.0 ± 2.0 beats/min (medium dose test) vs. 107.5 ± 7.5 beats/min (high dose test): P<0.05] and late phases [106.0 ± 6.7 beats/min (low dose test) vs. 84.0 ± 10.7 beats/min (medium dose test) vs. 117.5 ± 14.3 beats/min (high dose test): P<0.05] of reperfusion. The heart rate in the negative and the positive control groups increased from the equilibration values [150.0±13.7 beats/min (positive control) vs. 102.0 ± 13.9 beats/min (negative control)] in the early [146.0 ± 19.3 beats/min (positive control) vs. 135.0 ± 25.9 beats/min (negative control)] and late phases [175.0 ± 30.9 beats/min (positive control) vs. 205.0 ± 41.1 beats/min (negative control)] of reperfusion. A graphical representation of the heart rates in the equilibration, and reperfusion phases is shown in Figure 8.

**Figure 8.**
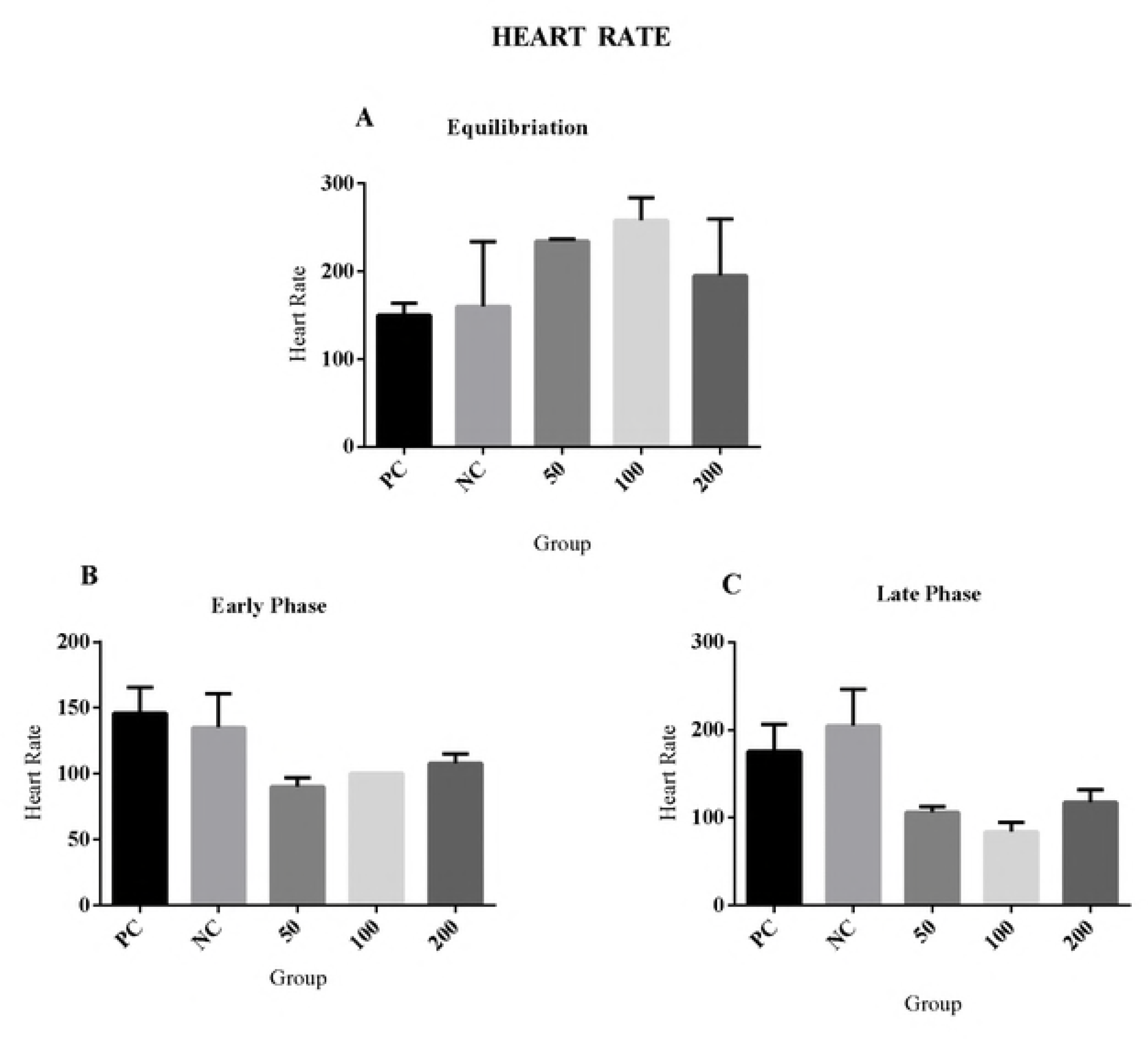
Heart rate; A) Equilibration heart rate values B) Early reperfusion phase heart rate C) Late reperfusion phase heart rate. LVDP (Left Ventricular Developed Pressure), PC (positive control), NC (Negative control)

### 3.3 Mechanism of action experiments

Co-administration of naloxone along the low dose test (50 mg) led to decreased +dp/dt_max_ recovery compared to treatment with 50 mg *S. coccinea* alone [0.09570±0.03294 (naloxone) vs.0.3868±0.1090(low dose test): P= 0.0338]. Similarly, naloxone led to a lower −dp/dt _min_ recovery compared to the 50 mg dose [0.7430±0.06739 (naloxone) vs.1.686±0.2693 (low dose test): P= 0.0179]. The graphical representation for the late recovery of the assessed indices of cardiac function when blockers were used is shown in Figure 9.

**Figure 9.**
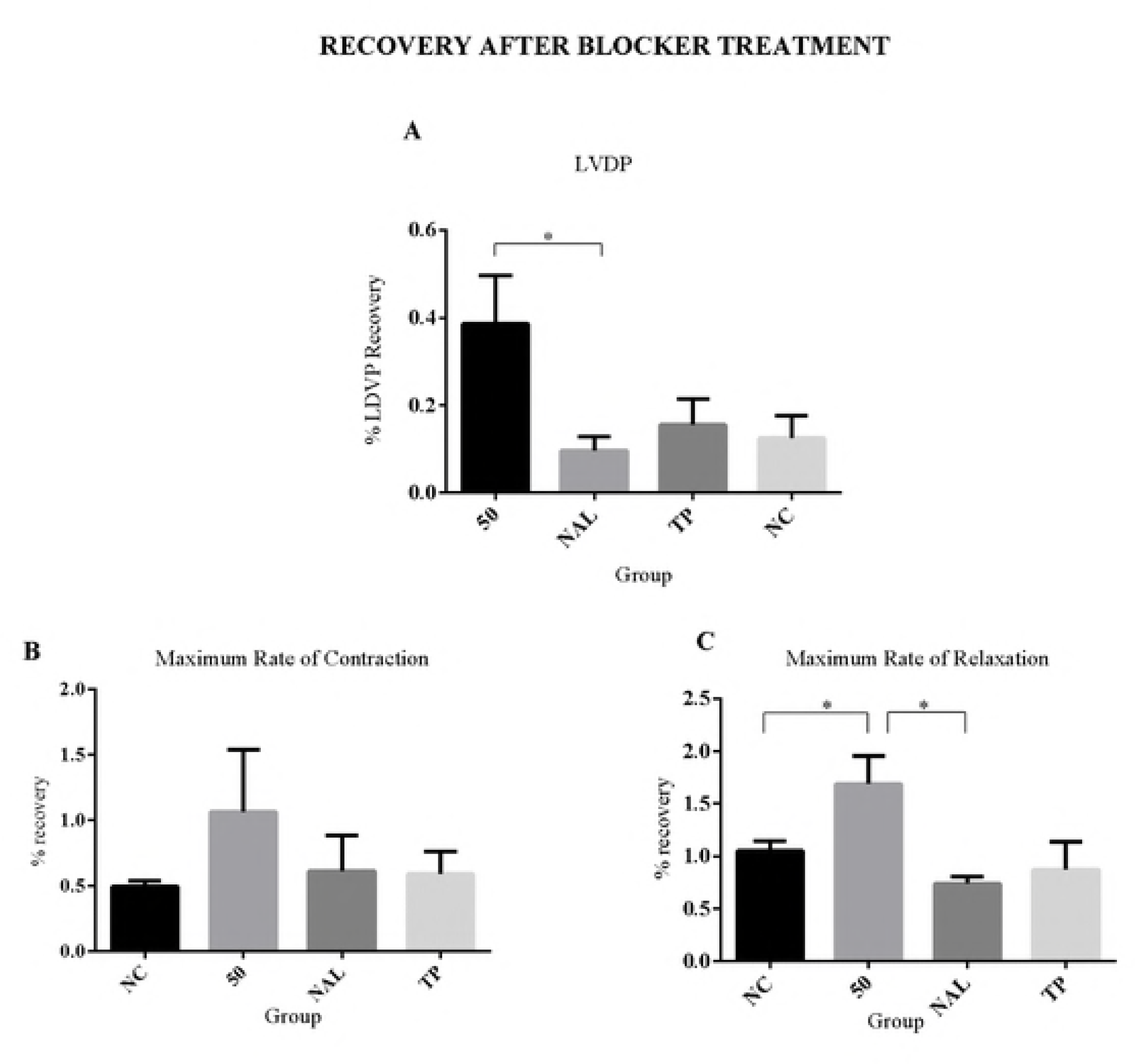
Indices of cardiac function. A) % recovery of Left ventricular developed pressure B)% recovery of maximum rate of contraction C) % recovery of maximum rate of relaxation. LVDP (Left Ventricular Developed Pressure), PC (positive control), NC (Negative control) KEY: * = P<0.05.

### 3.4 Dose-response analysis

The dose response curve for the cardioprotective effects of *S. coccinea* displayed a biphasic U-shaped pattern. As the dose was increased, the percentage recovery decreased up to a minimum point [x=153.2: y=1.0 (early phase) vs. x=173.97: y=12.49 (late phase)] beyond which the percentage recovery started to increase as shown in Figure 10.

**Figure 10.**
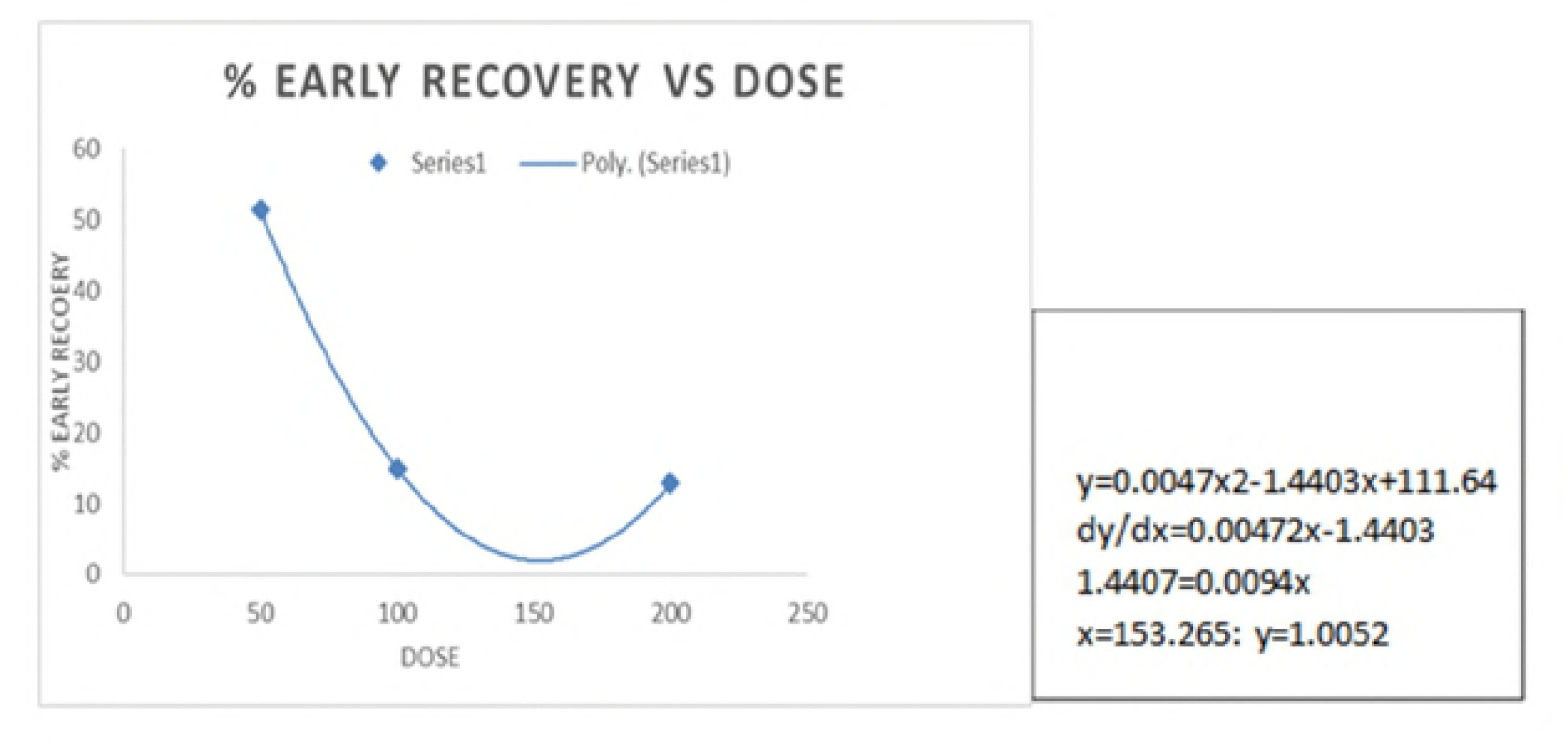
Dose response curve showing the percentage LVDP early reperfusion phase recovery after pharmacological post conditioning with the aqueous freeze dried extracts of *S. coccinea* following global ischemia.

## 4.0 Discussion

Ischemic heart disease (IHD) is among the leading causes of death and disability worldwide. Myocardial infarction induced by Ischemia/reperfusion (I/R) injury is one of the most severe manifestation of IHD, which makes its attenuation critical. The treatment of choice for myocardial ischemic injury is prompt and efficient reperfusion which paradoxically induces further cardiomyocyte damage known as reperfusion injury (9). Cardioprotection through pharmacological therapy in addition to early reperfusion is needed to further reduce the infarct size and reduce the mortality (10). Despite the burden caused by I/R injury there is still no effective therapy for preventing reperfusion injury which necessitates the search for more potent therapies (11).

Several salvia species are used as Danshen to treat heart diseases (5). However, the cardioprotective effects of *S. coccinea* and its possible mechanism of action have not yet been investigated.

Treatment with *S. coccinea* extract led to a significant dose-dependent recovery of the LVDP and +dp/dt _max_ after 30 minutes of global ischemia, with the highest recovery recorded at the low dose test (50 mg). Since LVDP and +dp/dt _max_ are measures of systolic function, this finding indicates that *S. coccinea* has the ability to prevent the systolic dysfunction that arises during I/R injury impairing the heart’s ability to build enough pressure at the required rate to pump blood commensurate to the body’s demands (12). These findings are in concurrence with those of a study that investigated the cardioprotective effects of another Salvia species: *S. miltiorhiza* (5). It is noteworthy that the percentage recovery in this study was higher compared to other cardioprotective herbal remedies (13).

The *S. coccinea* extract was shown to preserve the post-ischemic diastolic function in a dose dependent manner. The hearts treated with the herb had a higher percentage −dp/dt _min_ recovery compared to the control group hearts. Myocardial ischemia precipitates the development of diastolic dysfunction characterized by delayed relaxation, impaired filling and left ventricular stiffening (14). This finding is consistent with a previous study on other salvia species which preserved diastolic function (13). The finding was in contrast with other studies in which the −dp/dt _min_ recovered to a lower level than the equilibration value (15). Accordingly, the aqueous extract of *S. coccinea* may be more potent in abating the diastolic dysfunction that arises in I/R injury.

The *S. coccinea* extracts led to a significant decrease in heart rate at all doses in both the early and late phases of reperfusion in contrast to the control groups in which the heart rates were increased. Reperfusion following ischemia induces tachycardia which initiates life threatening ventricular arrhythmias (16). Low heart rates post-ischemia have thus been shown to be protective (17). These previous studies suggest that the heart rate lowering effects of *S. coccinea* might have been protective to the injured myocardium. This finding was however contradictory to *S. miltiorhiza* and its isolated components where the heart rate remained constant with no significant decrease (18).

This study also sought to assess the degree of cardioprotection by *S. coccinea* in the early and late phases of reperfusion. This is because most cardioprotective agents show potency only in the early phase (19). The events occurring in the early and late reperfusion phases differ and the cardioprotective potency of an agent in both phases depends on its ability to ameliorate the injurious events in both phases (20). Lethal reperfusion injury in the early phase is due to necrosis while that in the late phase is attributed to apoptosis (21). It is interesting to note that the cardioprotective effects of the *S. coccinea* extract were highly maintained in the early and late phases of reperfusion, a clear indication that the aqueous extracts may have the potential to prevent both necrosis and apoptosis.

*S. coccinea* was most potent at the 50 mg dose with the 100 mg and 200 mg providing cardioprotection to a lesser degree as indicated by the lower recovery of the LVDP, dp/dt _max_ and dp/dt _min_ in early and late reperfusion. This finding corroborates a study that investigated the cardioprotective effects of *S. miltiorhiza* where a lower dose, less than 100 mg was suggested as the 100 mg dose caused stiffening of the left ventricle and consequently diastolic dysfunction (13).

The dose response analysis of *S.coccinea* generated an inverted dose response curve with characterized by low dose inhibition and high dose stimulation of cardioprotection as opposed to the monophasic S-shape favored in pharmacological analysis (22). The probable molecular explanation for the inverted dose - response curve is biased agonism, a property of the ligand-receptor complex where a ligand favors one signaling pathway over another. β-arrestins usually associated with desensitization of the GPCRs, are involved in signaling (23) and have been shown to stimulate signal transduction pathways including MAPK, PKC, PI3K (24). Biochemical data shows that signaling mediated by β arrestins and G proteins have very distinct physiological consequences (25).

Pretreatment with naloxone at the onset of reperfusion completely abolished the cardioprotective effects of *S. coccinea*. This indicates that the cardioprotection by *S. coccinea* is mediated via the opioid pathway. Agonist binding to the opioid receptor usually inhibits adenylyl cyclase and activates phospholipase C signaling (26). Previous studies have shown that PLC signaling leads to the activation of the PKC and RISK kinases which are activated by salvianolic acid A and B which are phenolic compounds commonly found in salvia species (27). It is key to note that opioid receptors were among the first to portray biased agonism (24), a phenomenon that was observed in the dose response analysis of *S.coccinea*.

The synthesis and release of endogenous opioids into the circulation has been shown to increase in response to stressful stimuli including ischemia. Opioids mediate organ responses to stress by inducing hibernation and metabolic protection and are thus vital in enhancing the ability of the heart to withstand ischemia reperfusion injury (Akil et al., 1984). Myocardial binding studies have shown the presence of *kappa* and *delta* opioid receptors in ventricular myocytes. The capacity of the heart to synthesize all the three opioid peptides has been extensively verified experimentally (Krumins et al., 1985) (28) (29).

Pretreatment with theophylline partially abolished the cardio protection indicating that the adenosine pathway does not work in isolation to mediate the cardioprotection by *S. coccinea*. Adenosine receptors are known to interact with opioid receptors via receptor oligomerization (30). Adenosine is phosphorylated to AMP by adenosine kinase which is inhibited upon opioid receptor activation leading to a rise in adenosine levels (30). Consequently, it follows that, when theophylline blocks the adenosine receptors, only one pathway of cardioprotection is obstructed leaving the opioid pathway operational and hence the partially abolished cardioprotection.

Preliminary photochemistry analysis of *S.coccinea* confirmed the presence of saponins, sugars, phenols, tannins, amino groups, triterpenes, alkaloids and flavonoids (31). The pharmacological activities of the compounds from *S. coccinea* have not yet been fully elucidated and further studies would reveal the actual constituents that are involved in mediating the cardioprotective effects as observed in this present study.

Further work needs to be done to identify the active component of *S. coccinea* responsible for the cardioprotection observed. The cardioprotection experiments should also be performed in *in-vivo* models of ischemia.

## Conclusion

The freeze-dried extracts of *S. coccinea* possessed significant activity in the prevention of ischemia reperfusion injury. These actions were probably mediated by activation of the opioidergic system in the heart and lowering the heart rate through activation of the opioid pathway.

## Author Contributions

PWM, FOB and NMN designed the study and analyzed the experimental data. PWM and NMN wrote the paper. NMN did the experimental work. All authors read and approved the final manuscript.

## Acknowledgements

The authors wish to acknowledge the great technical assistance offered by Mr. David Kayaja Wafula and the technical staff in the department of medical physiology, the University of Nairobi in the execution of the study. The authors would also like to acknowledge the help from Mr. Patrick Mutiso Chalo of the University of Nairobi herbarium in the identification and collection of the plants.

## References

1. Lloyd-Jones D, Adams RJ, Brown TM, Carnethon M, Dai S, Simone GD, et al. Heart disease and stroke statistics - 2010 update: A report from the American heart association. Circulation. 2010 Feb 1;121(7):948–54.

2. Go AS, Mozaffarian D, Roger VL, Benjamin EJ, Berry JD, Blaha MJ, et al. Heart Disease and Stroke Statistics—2014 Update. Circulation. 2014 Jan 21;129(3):e28–292.

3. Heusch G, Libby P, Gersh B, Yellon D, Böhm M, Lopaschuk G, et al. Lancet Seminar: Cardiovascular Remodelling in Coronary Artery Disease and Heart Failure. Lancet. 2014 May 31;383(9932):1933–43.

4. Kin H, Zhao Z-Q, Sun H-Y, Wang N-P, Corvera JS, Halkos ME, et al. Postconditioning attenuates myocardial ischemia–reperfusion injury by inhibiting events in the early minutes of reperfusion. Cardiovasc Res. 2004 Apr 1;62(1):74–85.

5. Nie R, Xia R, Zhong X, Xia Z. Salvia miltiorrhiza treatment during early reperfusion reduced postischemic myocardial injury in the rat. Can J Physiol Pharmacol. 2007 Oct 1;85(10):1012–9.

6. Sage [Internet]. [cited 2018 May 27]. Available from: https://books.google.com/books/about/Sage.html?id=oHaMhvc_0dUC

7. Bilgin M, Sahin S, Dramur MU, Sevgili LM. Obtaining Scarlet Sage (salvia Coccinea) Extract Through Homogenizer- and Ultrasound-Assisted Extraction Methods. Chem Eng Commun. 2013 Sep 2;200(9):1197–209.

8. Mähler (Convenor) M, Berard M, Feinstein R, Gallagher A, Illgen-Wilcke B, Pritchett-Corning K, et al. FELASA recommendations for the health monitoring of mouse, rat, hamster, guinea pig and rabbit colonies in breeding and experimental units. Lab Anim. 2014 Jul 1;48(3):178–92.

9. Peuhkurinen K. [Myocardial reperfusion--a double-edged sword?]. Duodecim Laaketieteellinen Aikakauskirja. 1989;105(9):822–30.

10. Kloner RA, Schwartz Longacre L. State of the Science of Cardioprotection: Challenges and Opportunities— Proceedings of the 2010 NHLBI Workshop on Cardioprotection. J Cardiovasc Pharmacol Ther. 2011 Sep 1;16(3–4):223–32.

11. Fröhlich GM, Meier P, White SK, Yellon DM, Hausenloy DJ. Myocardial reperfusion injury: looking beyond primary PCI. Eur Heart J. 2013 Jun 14;34(23):1714–22.

12. McMurray JJV. Systolic Heart Failure. N Engl J Med. 2010 Jan 21;362(3):228–38.

13. Chang PN, Mao JC, Huang SH, Ning L, Wang ZJ, On T, et al. Analysis of Cardioprotective Effects Using Purified Salvia miltiorrhiza Extract on Isolated Rat Hearts. J Pharmacol Sci. 2006;101(3):245–9.

14. Mandinov L, Eberli FR, Seiler C, Hess OM. Diastolic heart failure. Cardiovasc Res. 2000 Mar 1;45(4):813–25.

15. Yu J, Wang L, Akinyi M, Li Y, Duan Z, Zhu Y, et al. Danshensu protects isolated heart against ischemia reperfusion injury through activation of Akt/ERK1/2/Nrf2 signaling. Int J Clin Exp Med. 2015 Sep 15;8(9):14793–804.

16. Lujan HL, DiCarlo SE. Reperfusion-induced sustained ventricular tachycardia, leading to ventricular fibrillation, in chronically instrumented, intact, conscious mice. Physiol Rep [Internet]. 2014 Jun 27 [cited 2018 May 27];2(6). Available from: https://physoc.onlinelibrary.wiley.com/doi/abs/10.14814/phy2.12057

17. Ischemia-induced and reperfusion-induced arrhythmias: importance of heart rate. Am J Physiol-Heart Circ Physiol [Internet]. [cited 2018 May 27]; Available from: https://www.physiology.org/doi/abs/10.1152/ajpheart.1989.256.1.H21

18. Qiao Z, Ma J, Liu H. Evaluation of the Antioxidant Potential of Salvia miltiorrhiza Ethanol Extract in a Rat Model of Ischemia-Reperfusion Injury. Molecules. 2011 Dec 2;16(12):10002–12.

19. Piper HM, Garcña-Dorado D, Ovize M. A fresh look at reperfusion injury. Cardiovasc Res. 1998 May 1;38(2):291–300.

20. Yellon DM, Hausenloy DJ. Myocardial Reperfusion Injury. N Engl J Med. 2007 Sep 13;357(11):1121–35.

21. Borutaite V, Jekabsone A, Morkuniene R, Brown GC. Inhibition of mitochondrial permeability transition prevents mitochondrial dysfunction, cytochrome c release and apoptosis induced by heart ischemia. J Mol Cell Cardiol. 2003 Apr 1;35(4):357–66.

22. Sonneborn JS. Mimetics of Hormetic Agents: Stress-Resistance Triggers. Dose-Response. 2010 Jan 1;8(1):dose-response.09-025.Sonneborn.

23. Biased agonism: An emerging paradigm in GPCR drug discovery. Bioorg Med Chem Lett. 2016 Jan 15;26(2):241–50.

24. Kelly E. Efficacy and ligand bias at the μ-opioid receptor. Br J Pharmacol. 2013 Jul 12;169(7):1430–46.

25. Rajagopal S, Rajagopal K, Lefkowitz RJ. Teaching old receptors new tricks: biasing seventransmembrane receptors. Nat Rev Drug Discov. 2010 May;9(5):373.

26. Maslov LN, Khaliulin I, Oeltgen PR, Naryzhnaya NV, Pei J-M, Brown SA, et al. Prospects for Creation of Cardioprotective and Antiarrhythmic Drugs Based on Opioid Receptor Agonists. Med Res Rev. 2016 May 16;36(5):871–923.

27. Inagaki K, Churchill E, Mochly-Rosen D. Epsilon protein kinase C as a potential therapeutic target for the ischemic heart. Cardiovasc Res. 2006 May 1;70(2):222–30.

28. Zhang Y, Irwin MG, Wong TM, Chen M, Cao C-M. Remifentanil Preconditioning Confers Cardioprotection via Cardiac κ- and δ-Opioid Receptors. Anesthesiol J Am Soc Anesthesiol. 2005 Feb 1;102(2):371–8.

29. Tanaka K, Kersten JR, Riess ML. Opioid-induced Cardioprotection. Curr Pharm Des. 2014;20(36):5696–705.

30. Peart JN, Gross GJ. Cardioprotection following adenosine kinase inhibition in rat hearts. Basic Res Cardiol. 2005 Jul 1;100(4):328–36.

31. Pérez S, C R de la, Canavaciolo G, L V, Marrero Delange D, Leyes R, et al. Análisis fitoquímico de la Salvia coccinea que crece en Cuba. Rev Cuba Plantas Med. 2011 Mar;16(1):54–9.

